# Global ocean resistome revealed: exploring Antibiotic Resistance Genes (ARGs) abundance and distribution on TARA oceans samples through machine learning tools

**DOI:** 10.1101/765446

**Authors:** Rafael R. C. Cuadrat, Maria Sorokina, Bruno G. Andrade, Tobias Goris, Alberto M. R. Dávila

**Affiliations:** Molecular Epidemiology Department, German Institute of Human Nutrition Potsdam-Rehbruecke - DIfE; Friedrich-Schiller University, Lessingstrasse 8, 07743 Jena, Germany; Embrapa Southeast Livestock - EMBRAPA; Department of Molecular Toxicology, Research Group Intestinal Microbiology, German Institute of Human Nutrition Potsdam-Rehbruecke - DIfE; Computational and Systems Biology Laboratory, Oswaldo Cruz Institute, FIOCRUZ. Av Brasil 4365, Rio de Janeiro, RJ, Brasil. 21040-900

**Keywords:** Beta-lactamase, NDM-1, marine metagenomics, Colistin, deep learning, OLS, multidrug resistance

## Abstract

The rise of antibiotic resistance (AR) in clinical settings is one of the biggest modern global public health concerns. Therefore, the understanding of AR mechanisms, evolution and global distribution is a priority due to its impact on the treatment course and patient survivability. Besides all efforts in the elucidation of AR mechanisms in clinical strains, little is known about its prevalence and evolution in environmental uncultivable microorganisms. In this study, 293 metagenomic from the TARA Oceans project were used to detect and quantify environmental antibiotic resistance genes (ARGs) using machine learning tools. After extensive manual curation, we show the global ocean ARG abundance, distribution, taxonomy, phylogeny and their potential to be horizontally transferred by plasmids or viruses and their correlation with environmental and geographical parameters. A total of 99,205 environmental ORFs were identified as potential ARGs. These ORFs belong to 560 ARG families that confer resistance to 26 antibiotic classes. 24,567 ORFs were found in contigs classified as plasmidial sequences, suggesting the importance of mobile genetic elements in the dynamics of ARGs transmission. Moreover, 4,804 contigs with more than 2 ARGs were found, including 2 plasmid-like contigs with 5 different ARGs, highlighting the potential presence of multi-resistant microorganisms in the natural ocean environment. This also raises the possibility of horizontal gene transfer (HGT) between clinical and natural environments. The abundance of ARGs showed different patterns of distribution, with some classes being significantly more abundant in coastal biomes. Finally, we identified ARGs conferring resistance to some of the most relevant clinical antibiotics, revealing the presence of 15 ARGs from the recently discovered MCR-1 family with high abundance on Polar Biomes. Of these, 5 were assigned to the genus *Psychrobacter*, an opportunistic pathogen that can cause fatal infections in humans. Our results are available on Zenodo in MySQL database dump format and all the code used for the analyses, including a Jupyter notebook can be accessed on GitHub (https://github.com/rcuadrat/ocean_resistome).

## Introduction

Antibiotic-resistant bacteria cause over 700,000 deaths per year, making them a global public health issue and an economic burden to the entire world and in particular in the developing countries. If the emergence of multi-resistant bacteria continues at the same rate, projections show that by 2050 they can cause 10 million deaths per year, which would outnumber deaths caused by cancer [1,2]. Despite its danger for human health, antibiotic resistance (AR) is a natural phenomenon and is one of the most common bacterial defence mechanisms. For example, the resistance to β-lactam antibiotics, conferred by beta-lactamase activity, is estimated to have emerged more than 1 billion years ago [3,4]. The collection of antibiotic resistance genes (ARGs) in a given environment, also called resistome, is a natural feature of microbial communities, being part of both inter- and intra-community communication and in the defence repertoires of organisms sharing the same biological niche [5,6].

Over the years, reservoirs of ARGs have been detected in different natural environments, such as oceans [7], lakes [8], rivers [9], remote pristine Antarctic soils [10] and impacted Arctic tundra wetlands [11]. Studies also show that anthropogenic activity (e.g. over-usage of antibiotics and their subsequent release via wastewaters into the environment) could lead to the spread of clinical important ARGs across natural environment [12,13]. Therefore, the investigation of the natural context of ARGs, their geographic distribution, dynamics and, in particular, their presence on horizontally transferable mobile genetic elements (MGEs), such as plasmids, transposons, and phages, is crucial to assess their potential to emerge and spread [14–16]. Due to modern advances in DNA sequencing and bioinformatics, it is now possible to study the presence and prevalence of ARGs in different environments. However, most of the published studies targeted only one or a few classes of ARGs and were limited to specific environments and geographic locations. The oceans cover around 70% of Earth’s surface, harbouring a big diversity of microscopic planktonic organisms forming a complex ecological network which is still under-studied [17,18]. To fix this, the number of ocean metagenomic projects stored in public databases have been growing, but the lack of related metadata made it difficult to conduct high-throughput gene screenings and correlations with environmental factors. Fortunately, the TARA oceans project [19] measured several marine environmental conditions, such as temperature, salinity, geographical location, pH, etc, across the globe and stored them as structured metadata. Together with the metagenome sequences [19], this allows the use of the machine and deep learning approaches to search for gene and species distribution and their correlation to environmental parameters.

In this study, we applied deepARG [20], a deep learning approach for ARG identification, to the 12 oceanic regions co-assembled TARA oceans contigs [21]. The results were then manually curated, a taxonomic classification was done, and analyses were performed for the gene abundance quantification together with association analyses between the quantification of ARGs and environmental parameters using Ordinary Least Squares (OLS) regression. We also explored the presence of ARGs in mobile genetic elements (MGEs) in TARA samples to investigate the potential of these oceanic environments to act as a reservoir of ARGs.

## Material and Methods

### Metagenomic data

A total of 12 co-assembled metagenomes (from different oceanic regions explored by the Tara Oceans expedition), with contigs larger than 1 kilobase were obtained from the dataset published in 2017 by Delmont et al. [22]. Raw reads of 378 shotgun sequencing runs of 243 samples were obtained from the EBI ENA database (https://www.ebi.ac.uk/ena) with accession numbers: PRJEB1787, PRJEB6606 and PRJEB4419.

Sample identifiers and metadata were obtained from Companion Tables Ocean Microbiome (EMBL) [23]. Tara Ocean samples were collected in seawater at different sites and depths and successively filtered using a single or a combination of membranes with pore sizes of 0.1□μm, 0.2□μm, 0.45□μm, 0.8□μm, 1.6□μm and 3□μm to retain different size fractions as viruses, giant viruses (“giruses”), prokaryotes (bacteria and archaea).

### Environmental ARG prediction

Open reading frame (ORF) prediction was performed on the 12 co-assembled metagenomes using the software MetaGeneMark v3.26 [24] with default parameters. The screening for ARGs was performed with DeepARG [20] on the extracted ORFs using gene models, classifying a gene as ARG if the probability was equal or greater than 0.8. In order to check if those contigs are plasmids or chromosomal sequences, contigs containing at least one putative ARG were analyzed with the tool PlasFlow 1.1 [25]. We also investigated the number and distribution of contigs with 2 or more putative ARGs to check for multiple resistance and/or whole ARG operons from environmental samples. ORFs of putative ARGs (and its respective contig) were submitted to Kaiju v1.6.2 [26] for taxonomic classification, with the option “run mode” set as “greedy”. Later, in order to check for misannotations and inconsistencies, we conducted a manual curation of each ARG among sequences from both deepARGdb [20] and those obtained from TARA contigs screening. Online BLASTp searches [27] were performed against the non-redundant protein database, with default parameters. Conserved domains (CDDs) and annotations in the source databases (ARDB, CARD and UniProt) were manually inspected.

### ARGs quantification and statistical tests on metagenomic samples

Environmental ARGs, identified after the manual curation, were used as a reference for the mapping of all the raw reads from the 378 metagenomic and 10 metatranscriptomic Tara Oceans samples, using BBMAP v37.90 [28]. The coverage, in terms of read counts and the abundance of each ARG was then calculated for each sample. Note that the abundance was calculated as Fragments Per Kilobase Million (FPKM). The Average Genome Size (AGS) and Genome Equivalents (GE) were estimated by the software MicrobeCensus v1.0.7 [29] in order to calculate Reads Per Kilo Genome equivalents (RPKG) as described by MicrobeCensus authors [29]. RPKG values for ORFs in each ARG family were summed for each sample. Environmental features, such as the sample depth, biogeographic biomes, ocean and sea regions and fractions (i.e. the concatenation of upper and lower filter size) were used for samples grouping and statistical tests. Pairwise Tukey HSD and multivariate linear regression using OLS models were conducted in Python 3.6 using the library ‘statsmodels’. The OLS was performed considering the following formula:

ARG_RPKG_ ∼ Marine provinces + Environmental Feature + Ocean sea regions + Fraction + Biogeographic biomes + Latitude + Longitude + NO2 + PO4 + NO2NO3 + SI + miTAGSILVATaxo Richness + miTAG_SILVA_Phylo_Diversity + miTAG_SILVA_Chao + miTAG_SILVA_ace + miTAG_SILVA_Shannon + OG_Shannon + OG_Richness + OG_Evenness + FC_heterotrophs_cells_mL + FC_autotrophs_cells_mL + FC_bacteria_cells_mL + FC_picoeukaryotes_cells_mL

where ARG_RPKG_ (the dependent variable) is the sum of RPKM of all ARGs in a given class and all the dependent variables are the selected environmental features. ANOVA was then conducted on the coefficients obtained from the OLS regression in order to infer the significance of a feature. A Python Jupyter notebook is provided with the code and the results for all the exploratory and statistical analyses [30].

### Phylogenetic analysis of environmental ARGs

Phylogenetic analyses were performed on the environmental genes identified as clinically relevant ARGs, such as the MCR-1 family and New Delhi metallo-beta-lactamase, for which reference clinical sequences were found in public databases, such as NCBI and deepARGdb. Multiple protein sequence alignments and phylogenetic trees were generated using the standard pipeline found on Phylogeny.fr [31]. Sequences were aligned using MUSCLE [32] in order to extract conserved blocks with gblocks [33] and phylogenetic trees generated with phyML [34], using “WAG” as substitution model and “alrt” as the statistical test.

### Database design and implementation

A manually curated MySQL database was created with the environmental ARGs described and all the subsequent analysis results. We provide the SQL dump at https://doi.org/10.5281/zenodo.3404245.

### Dash web application for data exploration and visualization

We developed a Python dashboard web application where the user can explore the results through interactive graphics (plotted with the plotly library). The app includes a geographical scatterplot, where it is possible to see the abundance of each ARG (or antibiotic class) selected by the user (RPKG) across all the samples in a world map; a boxplot, where the environmental feature can be selected in order to group the samples and compare the abundances; a barplot with taxonomic classification of the selected ARG (different taxonomic levels for the visualization can be selected); another scatterplot with marginal distribution plots and trend line (OLS), where on the axis X is represented the selected ARG and on the axis Y the user can select environmental variables (e.g. oxygen concentration, salinity, temperature, depth, etc.). In addition, a table containing information about each ORF is displayed. The additional information includes ORF id, contig id, antibiotic class, deepARG probability value, plasmid classification by PlasFlow, taxonomic classification by Kaiju (on the deepest level), if the ARG is expressed in at least one sample, other ARGs in the same contig and the total of ARGs in the contig. A link to download the multi-fasta file of the selected ARG is also provided. The application can be accessed at http://resistomedb.com/.

### Pipeline and code availability

The code of the complete pipeline (Figure 1) is in Bash and Python and it is available at the project repository on GitHub [35].

**Figure 1:**
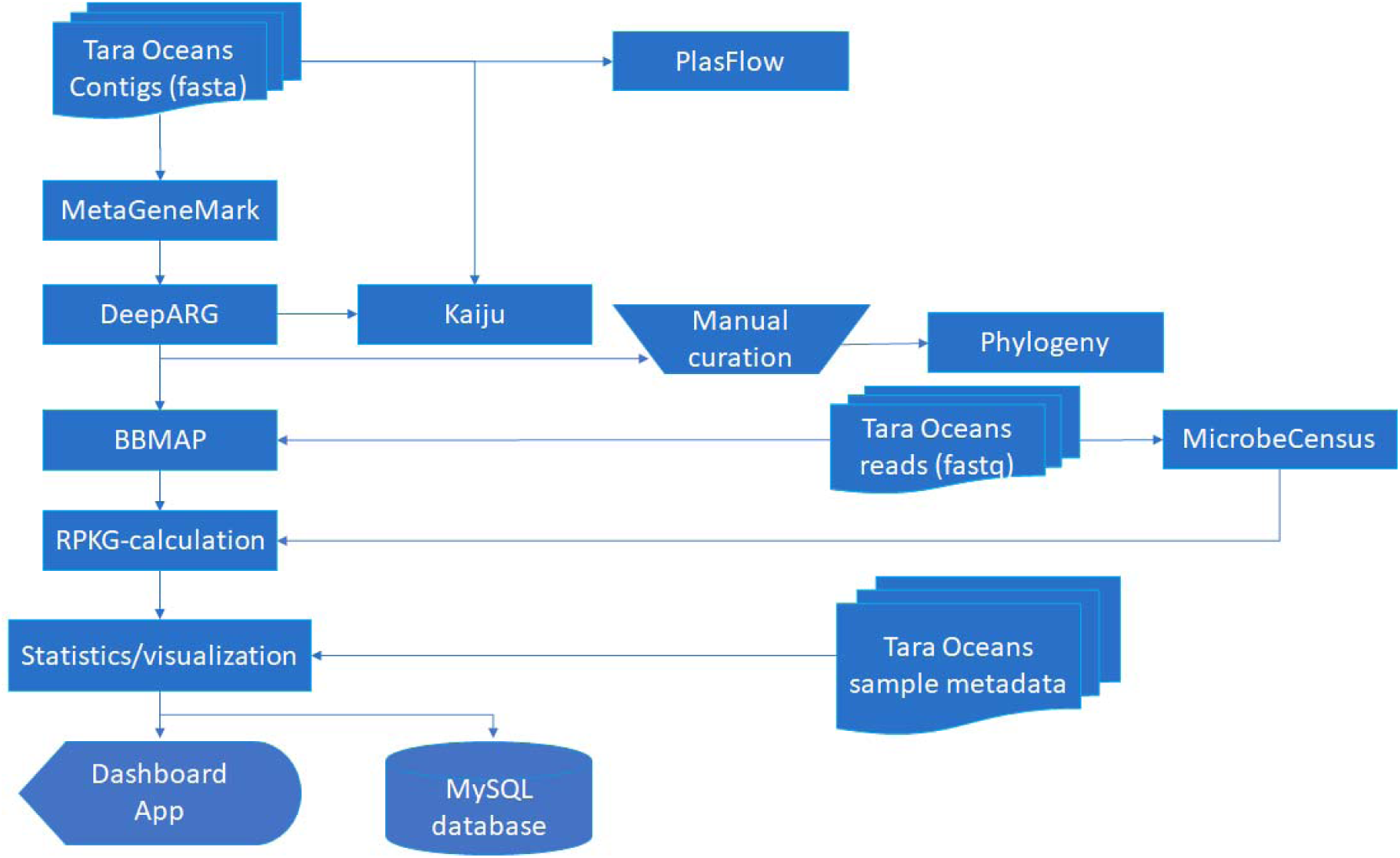
Flowchart used for ARG classification. The single steps and data used in the pipeline applied for the analyses presented in this work.

## Results and Discussion

### Environmental ARGs prediction and manual curation

A total of 41,249,791 ORFs were predicted by MetaGeneMark from 15,600,278 assembled contigs and used as input for ARG screening using the deepARG tool [11]. The usage of deepARG on the predicted Tara Oceans ORFs resulted in the classification of 116,425 ORFs (0.28%) as putative ARGs, belonging to 594 gene families and 28 ARG classes. The number of contigs, ORFs and putative ARGs of the metagenomic co-assemblies from each oceanic region can be found in Supplementary Table 1. Due to misannotations and misclassifications in the databases used for deepARG model training, it was necessary to conduct an extensive manual curation on the results. This curated dataset represents an important resource for further studies, including evolutionary and comparative studies.

The manual curation was performed to investigate the incidence of misannotated ARGs in different scenarios (Supplementary Table 2): (i) misannotated genes or gene families in the databases with low supportive evidence for ARG prediction; (ii) housekeeping genes that confer resistance only when specific mutations arise; (iii) housekeeping genes conferring resistance when overexpressed; (iv): regulatory sequences, responsible for ARG activation or overexpression of housekeeping genes (leading to a resistance phenotype); (v) sequences with both similarity to ARGs and non-ARGs, belonging to the same superfamily and/or sharing domains. All misannotated ARGs identified in scenario (i) were completely removed from our database and downstream analysis. A total of 34 ARG families was identified as misannotated or with low-quality annotation in the source database. For instance, the *msr*B gene encodes an ABC-F subfamily protein, that leads to erythromycin and streptogramin B resistance, but the fasta sequence in the database belongs to the *msrB* gene encoding methionine sulfoxide reductases B, not conferring antibiotic resistance. Another misannotated ARG family is *pat*A, an ABC transporter of *Streptococcus pneumoniae*, conferring resistance to fluoroquinolones, whose sequence is a putrescine aminotransferase (*pat*A) in the CARD database. A total of 99,205 ORFs remained after this step for non-quantitative analyses.

Putative ARGs identified on the scenarios (ii), (iii), (iv) and (v) were kept in the MySQL database for further studies but not used in the quantification and statistical analyses. The scenario (ii) includes the identification of 10 families of housekeeping genes and the corresponding mutations that could infer resistance. In scenario (iii), we identified 9 ARGs whose overexpression can lead to resistance. For scenario (iv) we identified 41 regulatory sequences that have been identified as responsible for ARG expression or over-expression of housekeeping genes that are leading to the resistance phenotype. For scenario (v) we identified 187 families that cannot be distinguished from non-ARGs by similarity alone. After the removal of these genes, a total of 13,163 ORFs (from the initial 116,425) belonging to 313/594 families, were retained for quantification and further analysis (Supplementary Table 2).

The most abundant ARGs of these 13,163 ORFs were QAC (multidrug efflux pumps named after their conferring resistance to quaternary ammonium compounds) with more than 3,000 overall occurrences, followed by TETB(60) with more than 2,000 occurrences (Figure 2). The latter is an ABC transporter that confers resistance to tetracycline and tigecycline identified by screening a human saliva metagenomic library [36]. The ORFs conferring resistance to tetracycline combined are the most widespread, with several TET and TETA classes accounting for approximately 4,000 occurrences. The main beta lactam-resistance conferring ARG was identified as K678_12262 with approximately 1,000 occurrences.

**Figure 2:**
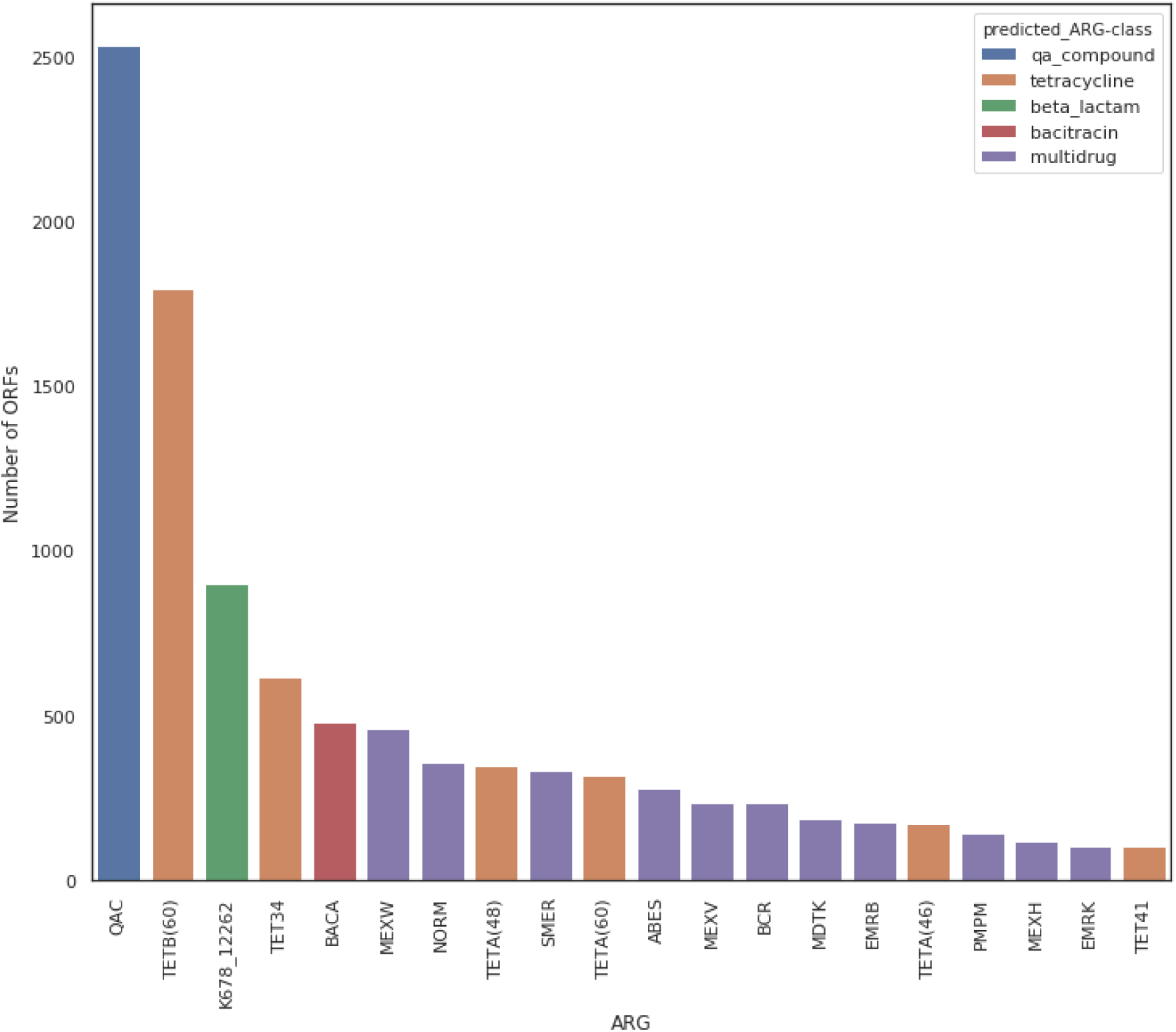
The 20 most abundant ARGs after manual curation. Number of ORFs detected in all metagenomes; the corresponding resistance to antibiotic classes is depicted in the upper right.

### Multiple ARGs in chromosomes and plasmids and taxonomic classification

We found a total of 24,567 putative ARGs (24.76% of the 99,205 ARGs excluding scenario (i)) present in contigs classified as plasmids. The presence of two or more ARGs in a single contig was also checked in order to identify possible multi-resistant organisms. The main objective of this analysis was to evaluate the potential of horizontal genetic transfer (HGT), as plasmids, bacteriophages, transposons, and extracellular DNA are the primary drivers of HGT. The occurrence of HGT of ARGs was already detected and characterized in clinical environments [37], in wastewater treatment plants (activated sludge) [14,38] and in fertilized soil [39], but still little is known about ARG HGT in aquatic environments, especially in open ocean regions. The presence of ARGs in phages and its potential HGT was described in some studies, for example, in a Mediterranean river [40], in pig faecal samples [15], in fresh-cut vegetables and in agricultural soil [16].

For this analysis, we only removed the ARGs from the scenario (i), misannotated sequences, because the presence of putative ARGs in the same contig and/or plasmid can give us additional functional evidence. We identified 4,063 contigs with multiple putative ARGs in contigs classified as chromosomes (up to 11 ARGs in the same contig), and 741 in contigs classified as a plasmid (up to 5 ARGs in the same contig), suggesting the presence of multi-resistant microorganisms in these environments (Table 1). In figure S1, we show the distribution of the ARGs in the 2 plasmids containing 5 ARGs each.

**Table 1:** Distribution of multiple ARGs in chromosome and plasmids. (classified by PlasFlow).

| Number of ARGs | in chromosome | in plasmid |
| --- | --- | --- |
| 2 | 3503 | 689 |
| 3 | 365 | 37 |
| 4 | 116 | 13 |
| 5 | 35 | 2 |
| 6 | 22 | 0 |
| 7 | 10 | 0 |
| 8 | 6 | 0 |
| 9 | 2 | 0 |
| 10 | 2 | 0 |
| 11 | 2 | 0 |

The taxonomic classification of putative ARGs performed using Kaiju [26], allowed us to classify 97,244 ARGs (98.02% of all ARGs excluding scenario (i)) up to some taxonomic level. Alphaproteobacteria (37,360 sequences) were revealed as the largest taxonomic unit. A total of 124 ARGs were classified as a virus. The most abundant taxonomic viral group was assigned to unclassified Prymnesiovirus (21 ARGs) and *Chrysochromulina ericina* virus (CeV) (19 ARGs). However, all the 124 viral ARGs are in scenario (v), and further investigations should be done in order to curate these sequences.

For the contigs containing multiple ARGs, the taxonomic classification of each ARG was checked together with a full contig sequence classification by kaiju to verify if the ARGs in the same contig belong to the same organism, giving us an indication of possible mis-assembly or HGT. One contig with 11 ARGs (TARA_ANW-k99_1343221) was classified as HGW-Alphaproteobacteria-12, while two of the single ARGs were classified as *Parvibaculum lavamentivorans*, an alphaproteobacterial species first isolated from activated sludge in Germany [41]. The other 9 ARGs, were classified as HGW-Alphaproteobacteria-3, HGW-Alphaproteobacteria-12, and as generic Alphaproteobacteria. A previous study showed the presence of ARGs in a strain of *Parvibaculum* from marine samples by functional metagenomics [7] which might hint to a broader ARG distribution among this clade. The other contig with 11 ARGs (TARA_ANE-k99_4428305) was classified (whole contig as well as the single ARGs) as *Micavibrio sp*., an obligately predatory bacterium exhibiting ‘vampire-like’ behaviour on gram-negative pathogens [42]. First isolated from wastewater samples, this genus has been considered as a potential new therapeutic approach against multi-resistant bacteria [43] including MCR-1 positive strains [43], due to the fact that no species from the genus *Micavibrio* was found to be pathogenic for humans [44]. However, if *Micavibrio* species contain indeed multiple ARGs, this would raise concerns about any clinical therapeutic approaches with these bacteria. One of the putative plasmids containing five ARGs (contig TARA_PSE-k99_4996023 on figure S1) showed taxonomic agreement between the classification of its single ARGs, all assigned to *Tistrella mobilis*. This species was firstly isolated from Thailand wastewater [45], and later another strain was isolated from the Red Sea [46]. The other contig classified as a plasmid with 5 ARGs was classified as *Halomonas desiderata*, a denitrifying bacterium isolated first from a municipal sewage treatment plant [47]. From the 5 ARGs in this contig, 2 confer resistance to trimethoprim (DFRE and DFRA3). Previous work showed that another bacteria from the same genus (*Halomonas marisflavi* type strain) is resistant to trimethoprim in vitro [48]. However, in the same study, *Halomonas desiderata* did not show resistance phenotype for any of the antibiotics tested. For both genera, *Tistrella* and *Halomonas*, mega plasmids were described [46,49].

### ARGs quantification on metagenome and metatranscriptome samples

The average genome size of Tara Oceans samples were calculated by MicrobeCensus v 1.0.7 [29] and then used to estimate RPKG, a normalization method for gene abundance in microbial studies used to avoid the bias of genome sequence coverage between samples populated with different genome sizes [29], which is expected due to filtration methods applied in the TARA study.

Since the Tara Oceans samples had different lower and upper thresholds of filtration size, we grouped samples by both thresholds to compare the AGS. MicrobeCensus uses a set of gene markers to infer the AGS, and since these gene markers are mainly present in cellular (not viral) genomes, the fractions enriched for virus and giant virus (< 0.22 um and 0.1-0.22 um) showed very biased and aberrant results for AGS, which was very high, up to 395.4 megabases; and genome equivalents. This is because the AGS values are inversely proportional to the number of reads mapping to the housekeeping gene markers, that are in very low abundance in virus-enriched samples. Based on that, we decided to remove virus-enriched fractions from the quantitative analysis, keeping 293 sample runs for downstream quantitative analyses. In order to compare the abundance of ARG classes across different marine regions, fractions and layers, we ran pairwise Tukey HSD tests on the ARG classes (the sum of RPKG for each class). For example, we wanted to investigate if the coastal biomes contained significantly more ARGs from any class than other more pristine biomes. The quinolone and bacitracin ARG classes were significantly more abundant in the coastal biome than in the westerlies biome (adjusted p-values 0.0169 and 0.0076, respectively). Furthermore, the fosmidomycin and quinolone ARGs were significantly (adjusted p-value 0.0014 and 0.0438, respectively) more abundant in the coastal biome than in the trades biome (Figure 3, Supplementary Table 2). The high abundance of ARGs of quinolone class along other ARG classes was previously reported on China’s coastal environment [50]. These results could indicate that specifically, this class of ARGs is under anthropogenic pressure, and future studies should be carried to investigate it in greater detail.

**Figure 3:**
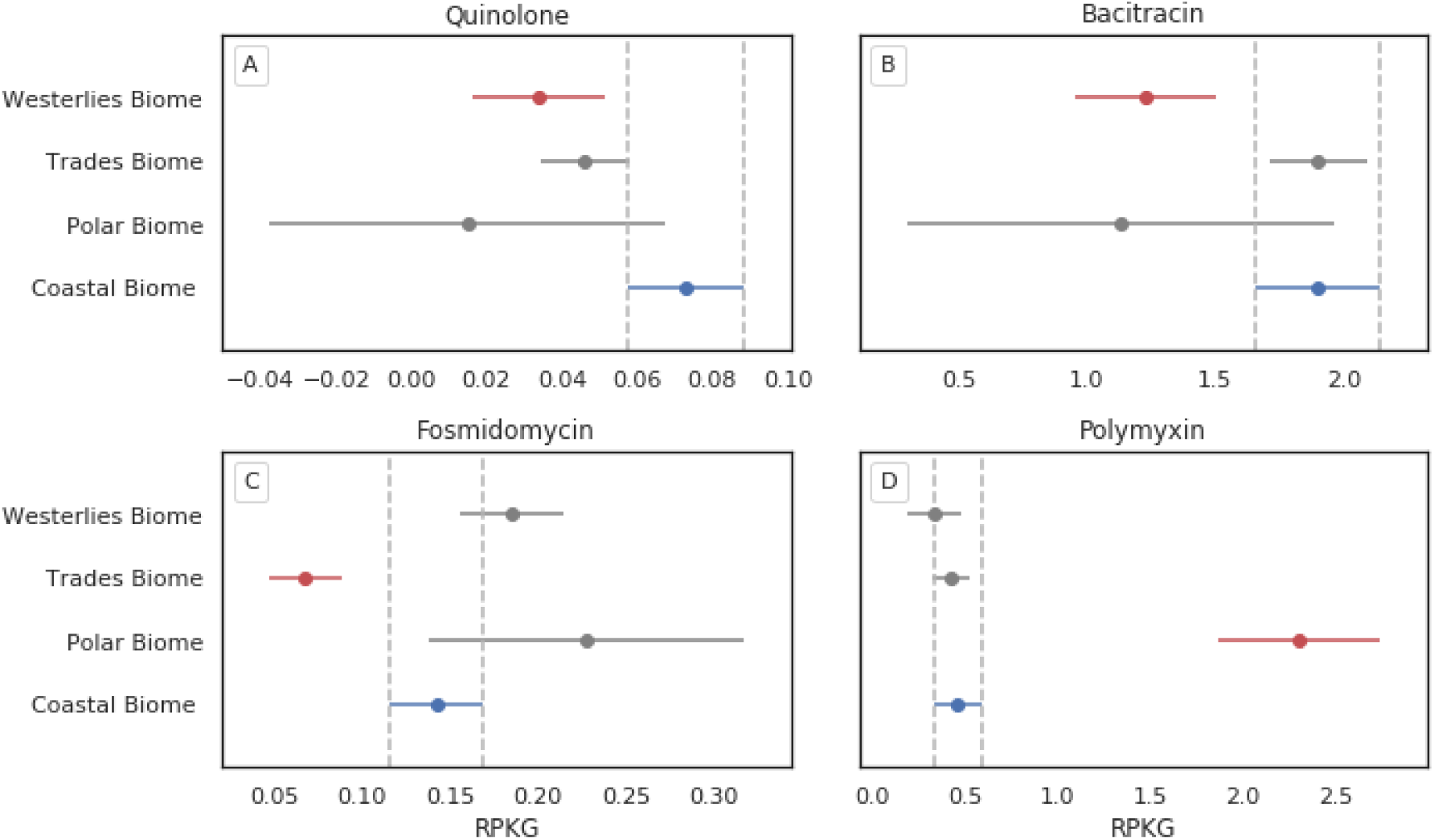
Significantly different abundances of ARG classes from Oceanic Biomes. Tukey HSD comparing the RPKG of ARG classes for 4 biomes of Tara Oceans study. A-RPKG for Quinolone ARGs; B-RPKG for Bacitracin ARGs; C-RPKG for Fosmidomycin ARGs; D-RPKG for Polymyxin ARGs. Reference for the test is in blue and in red the biome significantly different than the reference (p <0.05).

Surprisingly, the pristine polar biome showed significantly higher RPKG values for ARGs from the class Polymyxin than any other biome. Polymyxin B and E (also known as Colistins) are last-resort antibiotics used against gram-negative bacteria when modern antibiotics are ineffective, especially in cases of multiple drug-resistant *Pseudomonas aeruginosa* or carbapenemase-producing Enterobacteriaceae [51,52]. We discuss mobilized colistin resistance genes (MCRs) in greater detail in a separate section later in this manuscript.

When comparing abundances of classes on marine provinces, we found, for example, a significant difference (p < 0.05) of bleomycin class in 2 Indian provinces compared to most of the other provinces (Figure 4). Bleomycin resistance genes were previously reported to be in association with New Delhi metallo-β-lactamase (NDM-1) genes [53,54]. In this study, NDM-like genes (classified by deepARG as NDM-17 variant) were also found in greater abundance in India South Subtropical Gyre province. The first variant of NDM was identified in *Klebsiella pneumoniae* strain isolated from a Swedish patient who travelled to New Delhi, India [76], and it was spread globally in a few years and also in other species from the Enterobacteriaceae family, being classified as a potential worldwide public health problem [77]. This result raises concerns about the impact of antibiotics from the production of pharmaceuticals and wastewater in marine environments.

**Figure 4:**
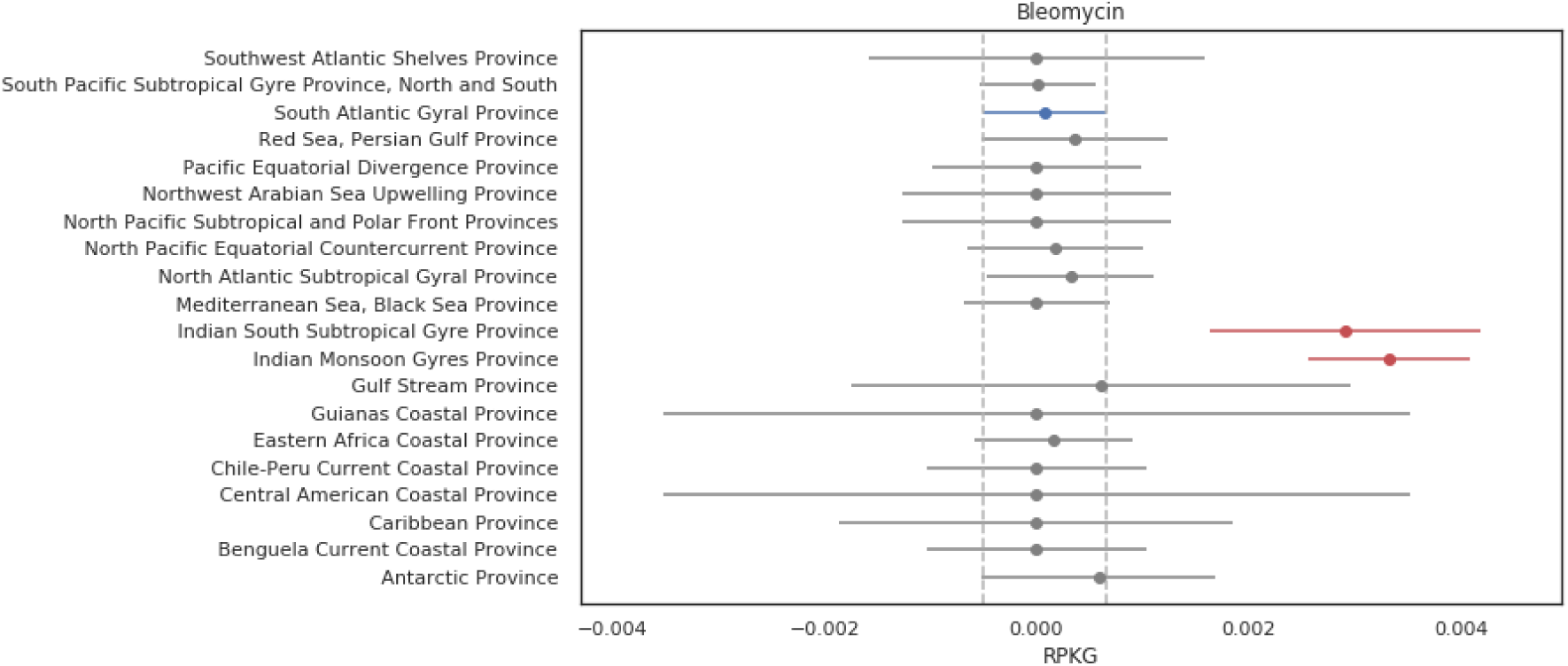
Bleomycin ARG abundance in marine provinces. Tukey HSD comparing the RPKG of ARGs from the class bleomycin. Reference for the test is in blue and in red the biome significantly different than the reference (p <0.05)

In addition, we wanted to check the influence of other environmental parameters on the abundance of ARG classes. Our OLS models showed mostly geographical parameters affecting the variance of ARGs, in agreement with our Tukey HSD tests. However, for some classes, the influence of non-geographical parameters (such as nutrient concentration) on the abundance of ARGs could be demonstrated. For example, for quinolone, the model shows inorganic phosphate (PO4)^3-^ inverted correlated (beta −0.13, p-value 0.0002), NO_2_NO_3_ (Nitrite+Nitrate concentration) direct correlated (beta 0.007, p-value 0.002). The R^2^ of this model is 0.57 and p-value 9.82E^-17^. On the other hand, PO4 is positively correlated with polymyxin and fosfomycin (p-values 1.918251E^-05^ and 1.651744E^-08^, respectively). The role of inorganic nutrients concentration in antibiotic resistance genes abundance is poorly understood and sometimes controversial, with some studies suggesting that high concentration of nutrients is negatively correlated with ARGs because in nutrient-rich environments competitive interactions are less important [55]. However, in wastewater treatments plants [56] and agricultural soil receiving dairy manure [57] the abundance of ARGs are increased. Further studies should be conducted to better understand the role of different nutrients on the abundance of ARGs of different classes in both pristine oligotrophic and impacted environments. The supplementary table 3 shows all significant results ANOVA test on the coefficients of OLS for each class.

### Mobilized colistin resistance genes (MCR-1) and other polymyxin resistance genes

Most mechanisms conferring resistance to Colistin are directed against modifications of the lipid A moiety of lipopolysaccharide (LPS), with the addition of l-ara4N and/or phosphoethanolamine (PEtN) to lipid A as the main mechanisms [58]. We found evidence for the occurrence of mobilized colistin resistance genes related to the recently discovered MCR-1 [59], which relies on the PEtN addition to lipid A. MCR-1 enzyme was described as 41% and 40% identical to the PEA transferases LptA and EptC, respectively, and sequence comparisons suggest that the active-site residues are conserved. However, until the discovery of the plasmid-borne mcr-1 in *E. coli* from pig [59], colistin resistance has always been linked to chromosomally encoded genes with few or no possibility of horizontal transfer. Further studies showed a high prevalence of the mcr-1 gene (e.g. 20% in animal-specific bacterial strains and 1% in human-specific bacterial strains in China) and the plasmid has been detected in several countries covering Europe, Asia, South America, North America and Africa [60–67]. Further MCR classes were described recently, with recent classification into 9 phylogenetically different classes MCR-1 to 9 [68,69]. In the present study, we found 15 ORFs classified as MCR-1 by deepARG, they were most abundant in the Atlantic Southwest Shelves Province followed by its adjacent region, Antarctic Province (Figure 5). However, the employed version of deepARG did not classify these sequences into the more recently described MCR-2 to 9. Therefore, we performed a phylogenetic analysis (Figure 6), which included sequences of different MCRs (MCR-1 to 5) and LptA (*eptA* as outgroup). The results suggested that 5 ORFs (from genus *Psychrobacter*, family *Moraxellaceae* [65]) are close to the MCR-1/2 clade with support value 1 (Figure 7). Members of the genus *Psychrobacter* were isolated from a wide range of habitats, including food, clinical samples, skin, gills and intestines of fish, seawater and Antarctic sea ice [67–71]. Importantly, at least 2 isolates from this genus were already reported to be resistant to Colistin (*Psychrobacter vallis sp. nov*. and *Psychrobacter aquaticus sp. nov*) both isolated from Antarctica [72]. Coincidently, the regions with greater RPKG mean for MCR-1 in our study were Southwest Atlantic and Antarctic Province. Our results support that *Psychrobacter* might be an ecological reservoir for transfer of PEtN transferases to other pathogens and further studies should be conducted to better understand the dynamics and evolution of ARGs in this genus. In addition, some species of this genus were also reported to cause opportunistic infections in humans, including at least one case reported to be associated with marine environment exposure [74] In this context it is therefore important to increase monitoring, by, for example, including screenings for those genes.

**Figure 5:**
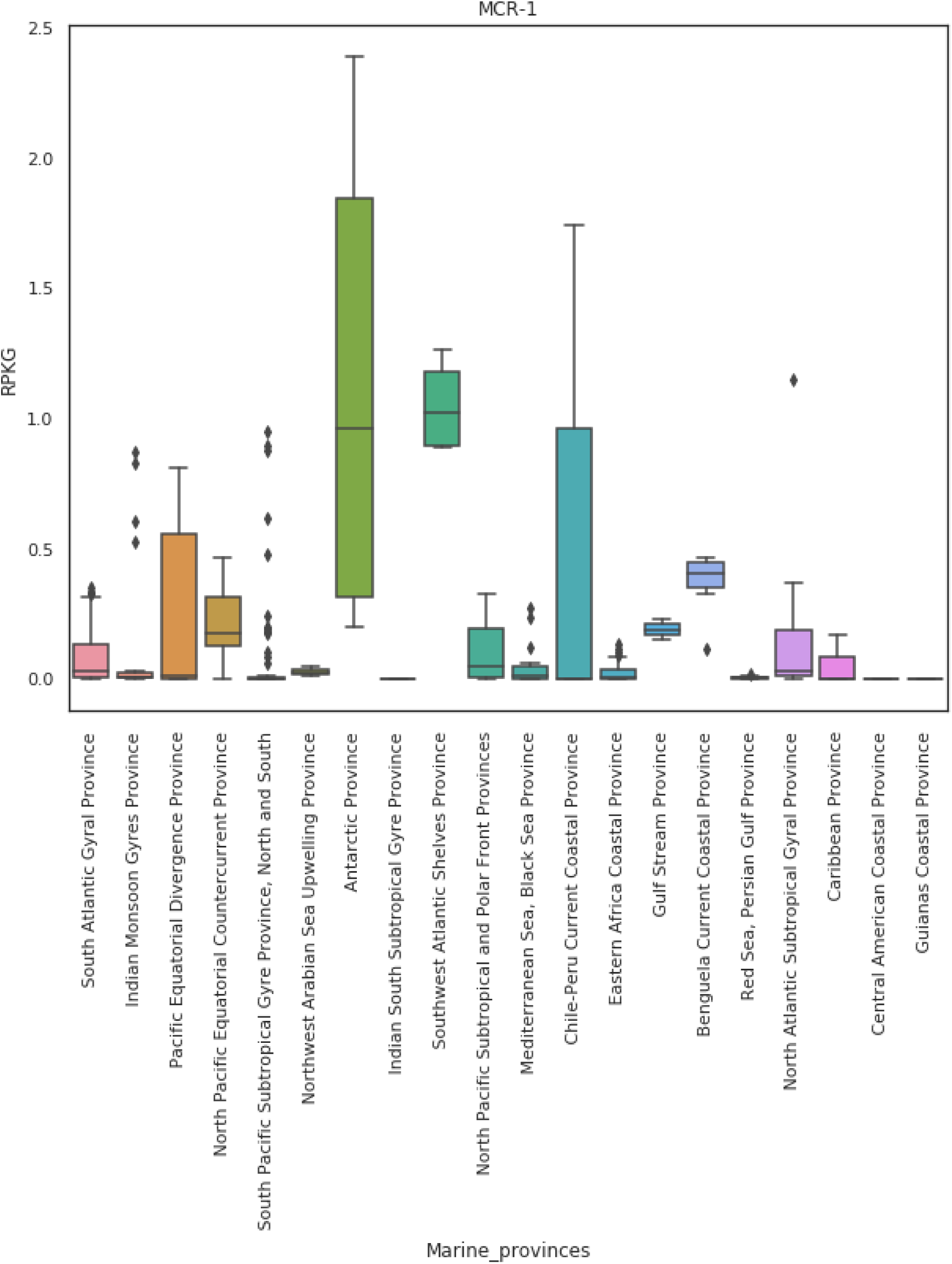
MCR-1 distribution on Tara Oceans marine provinces. The boxplot shows the sum of RPKG values for all MCR-1 ORFs.

**Figure 6:**
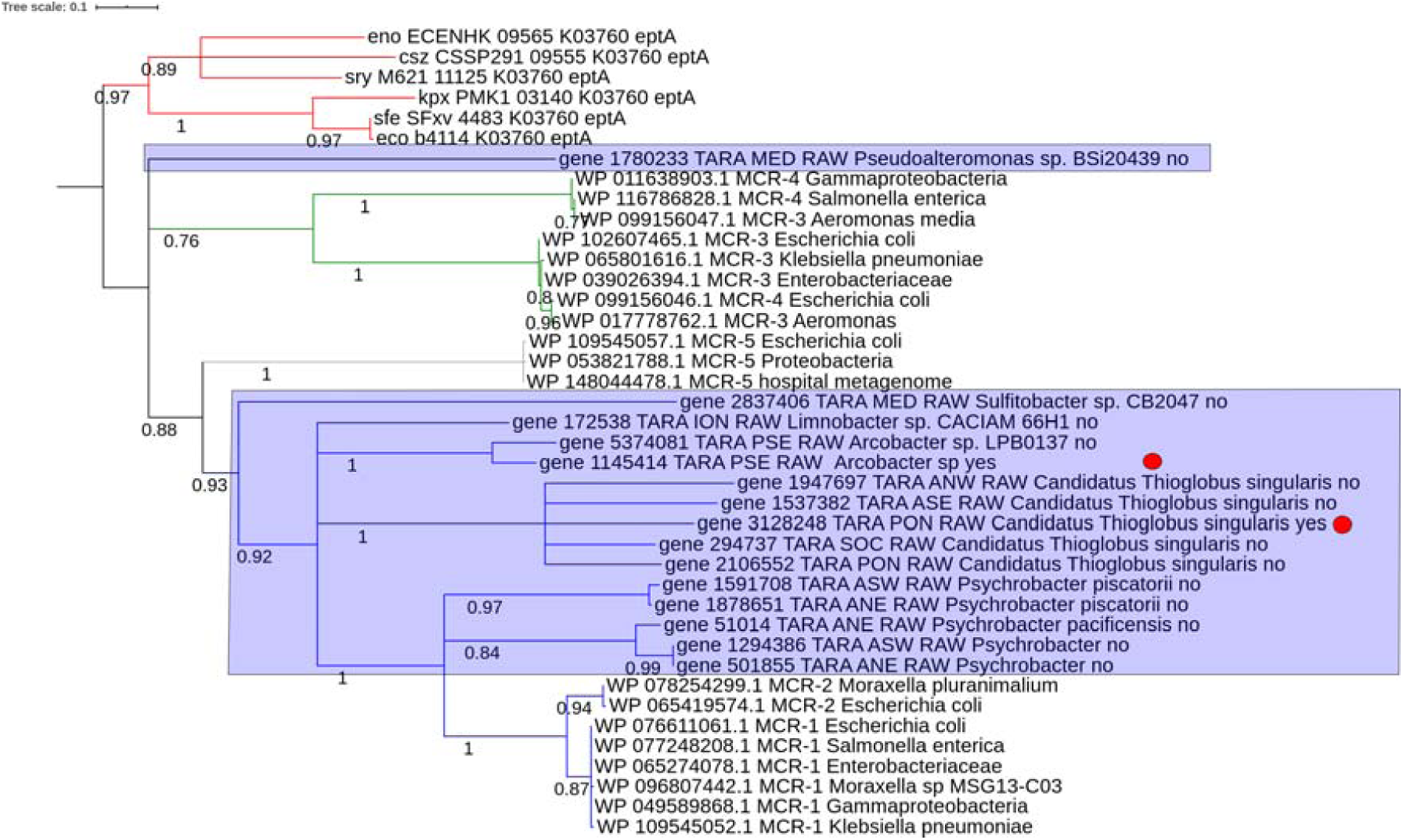
Phylogenetic tree of ocean metagenome MCR. The tree was inferred by standard pipeline from phylogeny.fr (phyML with “WAG” model and statistical test Alrt for support values). Sequences from NCBI for outgroup eptA and for clinical MCR-1 to MCR-5 were used in addition to the samples obtained from our results from Tara Oceans co-assemblies. The name of the Tara Oceans sequences displayed in the tree are defined with the id of sequence, co-assembly id, taxon name from Kaiju and yes/no for plasmid classification from PlasFlow. The blue rectangles are marking the TARA sequences. The blue clade is MCR-1/2 clade, the grey clade is MCR-5, the green clade is MCR-3/4 and the red clade is eptA clade (outgroup). The red circles are marking the sequences in contigs classified as plasmids by PlasFlow.

The residual MCR sequences, mostly belonging to the *Thioglobus* genus, were phylogenetically farther away from MCR-1/2 and might constitute new, distinct MCR classes. Important to note is that the phylogenetically very close relationship to MCR sequences does not prove the function as a colistin-resistant gene, which awaits further experiments to confirm this role.

For plasmid classification, we relied on the results of PlasFlow, which only classified two of the sequences classified located on a plasmid. This can be explained by the small size of many contigs (with eight of them smaller than 3 kb). Additionally, a false-negative result from PlasFlow could be a result of a re-integration of plasmidial sequences into the chromosome - or that the here detected MCR genes may constitute an ancestor of the plasmidial *E. coli* MCR sequences, as was already suggested for *Moraxella* species [66]. The two ARGs classified to be located on a plasmid are detected in contigs with a size of 2 kb and 38 kb. The former, classified as belonging to a *Thioglobus* species, is difficult to be validated as a plasmidial sequence due to its small size. The latter could be classified as a sequence of a *Poseidonibacter* species, a marine group of bacteria recently reclassified from the Arcobacter genus which contains several pathogenic bacteria [70]. A toxin-antitoxin system is encoded two ORFs upstream the MCR gene, which might be an indication for a plasmidial location. However, no further genes usually located on *Arcobacter* spp. plasmids [71] were found on this contig, hampering its definite classification as a plasmidial mcr. That said, various mobile element genes located on this contig (Figure 7) strengthen the assumption that this contig is related to a mobile genetic region. Interesting is the unusual synteny of MCR, PAP2 and a downstream encoded DAGK, of which the latter only appears in MCR-3 genetic environments [72]. Related genes (amino acid sequence identity of about 70%) with a conserved gene synteny are found in various Arcobacter species (Fig. 6). Several Arcobacter species which do contain a similar mcr gene were susceptible to colistin treatment (REF), arguing against this gene conferring colistin resistance. Further research would be necessary to confirm or refute colistin resistance in marine Poseidonibacter [73].

**Fig 7:**
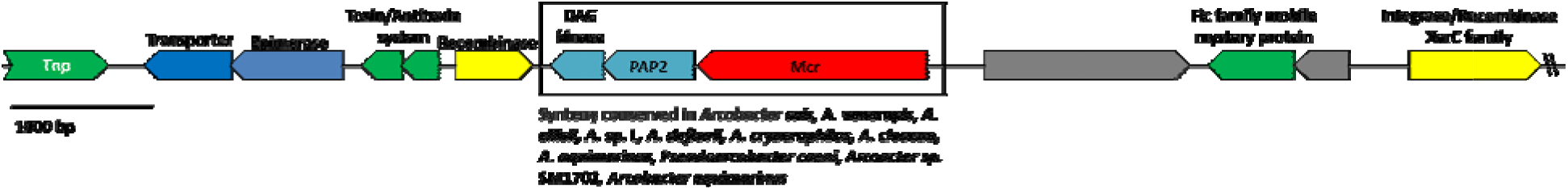
Genomic context of the mcr gene of contig TARA_PSE_k99_4834589. This contig was classified to be plasmidial by PlasFlow. Depicted are the first 13 ORFs from 28, showing MCR-1 and surrounding genes and including the mobile element related genes. Tnp - Transposase, DAG - Diacylglycerol, PAP2 - phosphatase PAP2 family protein, Mcr - mobilized colistin resistance protein. Colour code: green - mobile element related gene, blue - Other/metabolic genes, yellow - DNA-related gene, light blue - Mcr-accessory genes, red - Mcr gene, grey - hypothetical protein. Annotations from MetaGeneMark were manually refined using the conserved domains database and blastp against the SwissProt database. Taxonomy of *Arcobacter* species is stated as currently in the Genbank database.

The presence of MCR-related genes in both Antarctic and adjacent regions can also raise concerns about gene flow due to ice melting, a problem already discussed previously for other ARGs [73].

## Conclusions

Our study uncovers the diversity and abundance of ARGs in the global ocean metagenome, conferring probably resistance to 26 classes of antibiotics. The extensive analysis leads to a detailed taxonomic classification and varying ARG abundance in different biomes. Potential horizontal gene transfer leads to the spreading of some ARGs, resulting also in putative multi-resistant strains. For a limited number of ARGs, gene transcription could be shown. Our study also exposes the importance of monitoring coastal water for anthropogenic impact, since the inflow of antibiotics by e.g. wastewater might strengthen (by selective pressure) the antibiotic resistance development by microorganisms. Antarctic soil and ice might be a huge potential reservoir for ARGs, and the discussion about ARG distribution should not neglect the future impact of this reservoir under the influence of climate change. This study could also bear an impact on investigations dealing with the evolutionary history of ARGs, with the here presented genes as ancestors of common ARGs in clinically relevant strains. Last but not least, the combination of multiple modern machine learning tools and other open-source data science libraries such as Dash and Plotly produced a valuable resource for the scientific community working on further studies on antibiotic resistance in different environments.

## Acknowledgements

We thank Jorge Boucas and the Bioinformatics Core facility of Max Planck Institute of Biology of Ageing, for the use of the computational resources (HPC cluster) and the fruitful discussions in the initial analysis of this work.

## Competing Interests

The authors declare no competing interests.

## Notes

#### Summary of Updates

Improved discussion and figures. Included dashboard web application

https://zenodo.org/record/3404245#.XXgiaygzYuU

http://resistomedb.com/

