## Supplementary material for "Global ocean resistome revealed: exploring Antibiotic Resistance Genes (ARGs) abundance and distribution on TARA oceans samples through machine learning tools": SI

Supplementary Table 1: Number of contigs, ORFs and putative ARGs for each oceanic region (metagenomic co-assembly)

| **Sample** | **Region** | **Contigs** | **ORFs** | **#ARG** |
| --- | --- | --- | --- | --- |
| TARA_ANE_RAW | Atlantic North East | 1382239 | 3686619 | 11283 |
| TARA_ANW_RAW | Atlantic North West | 1267057 | 3308183 | 9994 |
| TARA_ASE_RAW | Atlantic South East | 766472 | 1842937 | 4955 |
| TARA_ASW_RAW | Atlantic South West | 989154 | 2502625 | 6902 |
| TARA_ION_RAW | Indian Ocean North | 1608737 | 4257000 | 9830 |
| TARA_IOS_RAW | Indian Ocean South | 1475271 | 3736229 | 9909 |
| TARA_MED_RAW | Mediterranean | 1146485 | 3344341 | 10160 |
| TARA_PON_RAW | Pacific Ocean North | 1663221 | 4368436 | 11814 |
| TARA_PSE_RAW | Pacific South East | 2722083 | 7155344 | 20587 |
| TARA_PSW_RAW | Pacific South West | 1124813 | 3080817 | 9224 |
| TARA_RED_RAW | Red Sea | 1039053 | 2937951 | 8924 |
| TARA_SOC_RAW | Southern Ocean | 415693 | 1029309 | 2843 |
| Total |  | 15,600,278 | 41,249,791 | 116,425 |

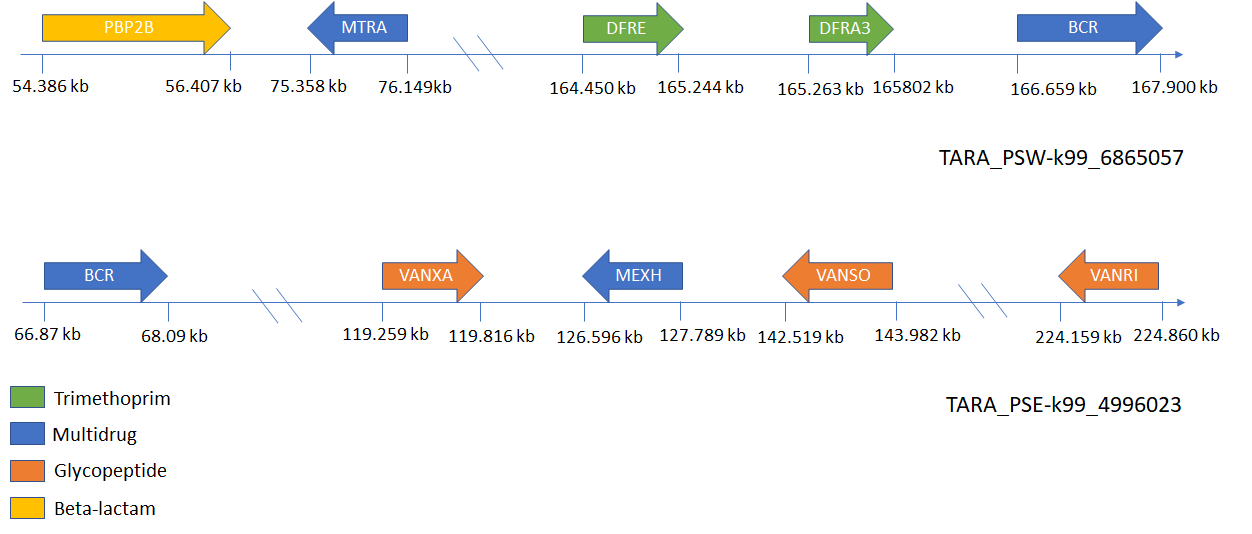

**Figure S1: ARGs distribution in the 2 plasmids showing 5 ARGs each.** The size of genes and distances are not scaled. **PBP2B**: methicillin-resistant PBP2; **MTRA**: transcriptional activator of the MtrCDE multidrug efflux pump; **DFRE**: dihydrofolate reductase; **DFRA3**: integron-encoded dihydrofolate reductase; **BCR**: Bicyclomycin resistance protein; **VANXA**: variant of VANX D,D-dipeptidase; **MEXH**: membrane fusion protein of the efflux complex MexGHI-OpmD; **VANSO**: variant of VANS, required for high-level transcription of other van glycopeptide resistance genes; **VANRI**: regulatory transcriptional activator in the VanSR regulator within the VanI glycopeptide resistance gene cluster.
